# Disrupted controlling mechanism of salience network on default-mode network and central-executive network in schizophrenia

**DOI:** 10.1101/2021.12.03.471183

**Authors:** Ganesh B. Chand, Deepa S. Thakuri, Bhavin Soni

## Abstract

Neuroimaging studies suggest that the human brain consists of intrinsically organized large-scale neural networks. Among those networks, the interplay among default-mode network (DMN), salience network (SN), and central-executive network (CEN)has been widely employed to understand the functional interaction patterns in health and diseases. This triple network model suggests that SN causally controls DMN and CEN in healthy individuals. This interaction is often referred to as the dynamic controlling mechanism of SN. However, such interactions are not well understood in individuals with schizophrenia. In this study, we leveraged resting state functional magnetic resonance imaging (fMRI) data of schizophrenia (n = 67) and healthy controls (n = 81) to evaluate the functional interactions among DMN, SN, and CEN using dynamical causal modeling. In healthy controls, our analyses replicated previous findings that SN regulates DMN and CEN activities (Mann-Whitney U test; p < 10^−8^). In schizophrenia, however, our analyses revealed the disrupted SN-based controlling mechanism on DMN and CEN (Mann-Whitney U test; p < 10^−16^). These results indicate that the disrupted controlling mechanism of SN on two other neural networks may be a candidate neuroimaging phenotype in schizophrenia.

## 1. Introduction

Schizophrenia is a severe neuropsychiatric condition currently affecting ~3 million people in the United States and ~7.8 billion people worldwide and therefore it has significant physical, emotional, and financial burden (Chong et al., 2016; Cloutier et al., 2016; Insel and Cuthbert, 2015; Kapur et al., 2012; McCutcheon et al., 2020). Despite extensive efforts, the underlying functional mechanisms of schizophrenia are not well understood. Previous studies suggest that normal cognitive functions involve the coordinated activity of intrinsically organized large-scale neural networks (Deco et al., 2011; Power et al., 2011). The three neural networks—salience network (SN), default-mode network (DMN), and central-executive network (CEN)—have been proposed to study the functional interactions among these networks in health and diseases (Chand and Dhamala, 2016a; Menon, 2011). SN, which primarily consists of the anterior insula and dorsal anterior cingulate cortex, responds to behaviorally salient events (Chand and Dhamala, 2016b; Seeley et al., 2007), and it plays a crucial role in dynamic cognitive activities, notably in initiating cognitive control, implementing task sets, and coordinating behavioral responses (Chand and Dhamala, 2017; Dosenbach et al., 2006; Menon and Uddin, 2010). DMN, which encompasses the posterior cingulate cortex and ventromedial prefrontal cortex, is involved in internally-oriented cognitive activities including autobiographical, self-monitoring, and social functions (Anticevic et al., 2012; Buckner et al., 2008). CEN, which comprises of the posterior parietal cortex and dorsolateral prefrontal cortex (DLPFC), is responsible for higher-level cognitive functions, including the control of attention, working memory, and decision-making (Bressler and Menon, 2010; Chand et al., 2016). Blood-oxygen-level-dependent (BOLD) fMRI investigations in healthy younger and elder populations have consistently illustrated that SN modulates the activity of DMN and CEN (Chand and Dhamala, 2016a; Chand et al., 2017b; Goulden et al., 2014; Sridharan et al., 2008; Wu et al., 2016). This SN-based modulation on DMN and CEN is also known as the dynamic controlling property of SN. The inter-connection patterns among these three networks and their coordination are not well understood in individuals with schizophrenia.

BOLD fMRI studies have previously defined a set of distinct large-scale neural networks related to active and passive cognitive states. Normal cognition involves the coordinated activity of large-scale neural networks; however, the way they interact is still unclear. Prior literature has investigated the coordinated activity among those networks and one of the theories argues that SN regulates neural activities of DMN and CEN in healthy controls (Chand and Dhamala, 2016a; Goulden et al., 2014; Menon, 2011; Sridharan et al., 2008; Wu et al., 2016). This SN-based control mechanism has been argued to be crucial for cognitive maintenance, including both resting state and tasks, in healthy controls. This mechanism has been formerly explained in terms of the unique cyto-architecture of SN nodes—anterior insula and dorsal anterior cingulate—that only these brain regions contain a special type of neurons called ‘von Economo neurons (VENs)’ (Allman et al., 2005). These VENs facilitate the rapid information processing within SN nodes and with other networks, including DMN and CEN. Prior studies further argue that there is an impairment of highly sensitive VENs in SN nodes with neuropsychiatric conditions, including schizophrenia (Butti et al., 2013). There are also reports of maximal gray matter reductions in SN nodes, especially insula (Gupta et al., 2015)—a key node of SN—of schizophrenia. However, SN-based dynamic coordinating property on DMN and CEN remains to be evaluated in schizophrenia. We hypothesized that SN-based modulations will decline in individuals with schizophrenia compared to healthy controls.

## 2. Materials and methods

### 2.1. Participants

In this study, data of 67 chronic schizophrenia patients and 81 healthy controls with an age range of 18-66 years were used. These data were assessed from a publicly available dataset (Aine et al., 2017; Cetin et al., 2014). Informed consent was obtained from each subject according to the local institutional guidelines for human subjects. Anonymized imaging data were shared by the SchizConnect (http://schizconnect.org) following the data request procedure. Schizophrenia patients were confirmed by completing the Structured Clinical Interview for DSM-IV Axis I Disorders. Schizophrenia patients with a history of neurological disorders, such as head trauma, mental retardation, or history of active substance abuse (except for nicotine) within a past year were excluded. Schizophrenia patients had a negative toxicology screen for drugs of abuse. Age- and sex-matched healthy controls were recruited and they completed the Structured Clinical Interview for DSM-IV Axis I Disorders—Non-Patient Edition to rule out Axis I conditions. Detailed inclusion and exclusion criteria are described in previous studies (Aine et al., 2017; Cetin et al., 2014).

### 2.2. Data acquisition

The Siemens 3T TIM Trio scanner with a 12-channel head coil was used for brain imaging. High resolution T1-weighted images were acquired with a five-echo multi-echo MPRAGE sequence using the following parameters: TE (echo times) = 1.64, 3.5, 5.36, 7.22, 9.08 ms, TR (repetition time) = 2.53 s, TI (inversion time) = 1.2 s, flip angle = 7 degrees, number of excitation = 1, slice thickness = 1 mm, FOV (field of view) = 256 mm, and resolution = 256 × 256. Resting state BOLD fMRI scans were collected using a single-shot gradient echo planar sequence with lipid suppression using the following parameters: TR = 2000 ms, TE = 29 ms, flip angle = 75 degrees, FOV = 240 mm, matrix size = 64 × 64, 33 slices, and voxel size: 3.75 × 3.75 × 4.55 mm.

### 2.3. Image Preprocessing

Resting state BOLD fMRI and T1-weighted anatomical images were preprocessed using SPM12 (Wellcome Trust Centre for Neuroimaging, UK;www.fil.ion.ucl.ac.uk/spm/software/spm12). The preprocessing steps included motion correction, co-registration to individual anatomical images, normalization to the Montreal Neurological Institute (MNI) space, and finally spatial smoothing of the normalized images with a 6 mm isotropic Gaussian kernel.

### 2.4. Independent component analysis and dynamical causal modeling

A constraint independent component analysis (ICA) was applied to preprocessed resting state BOLD fMRI data to extract the temporal signals of each network. ICA is a widely used method for functional brain network activities (Damoiseaux et al., 2012) and a spatially constrained ICA (Lin et al., 2010) overcomes the difficulties in identifying components of interest and in determining the optimum number of components over conventional ICA. Constrained ICA has been highlighted as a useful tool, particularly if one is interested in specific brain areas or network(s). Group ICA of fMRI Toolbox (GIFT; http://mialab.mrn.org/software/gift)was used to compute ICA components. Templates of each network (DMN, SN, and CEN) reported in previous study (Shirer et al., 2012) were used and the network-specific ICA components were computed. This approach renders better representation of networks compared to choosing an average or first eigen-variate of a template (Chand et al., 2018; Craddock et al., 2012; Goulden et al., 2014; Shirer et al., 2012; Smith et al., 2011).

For subsequent dynamical causal modeling (DCM) analysis, time courses (components) produced by constrained ICA were used. DCM assesses the directed connectivity between different brain areas or networks. In DCM, the hypothesized possible model architectures are specified and a Bayesian model selection (BMS) is then used to infer the model that best fits measured data (Stephan et al., 2010). DCM facilitates the implementation of random effects in model selection (Stephan et al., 2009) and allows the specification of nonlinear modulations to investigate how a neural network influences connection strengths between other neural networks (Li et al., 2011; Stephan et al., 2008). The stochastic model accounts for noise more accurately and therefore allows the application of DCM to the resting state fMRI (Daunizeau et al., 2012; Friston et al., 2014). DCM analyses were performed using SPM12 (Wellcome Trust Centre for Neuroimaging, UK; www.fil.ion.ucl.ac.uk/spm/software/spm12). We specified three models for each subject with full intrinsic connections (Friston et al., 2003; Friston et al., 2014) between the DMN, SN, and CEN consistent with our previous studies (Chand et al., 2017a, b). Model 1 specifies nonlinear modulations of DMN on the reciprocal connections between SN and CEN. Likewise, model 2 specifies nonlinear modulations of SN on the reciprocal connections between DMN and CEN. Lastly, model 3 specifies nonlinear modulations of CEN on the reciprocal connections between DMN and SN. The non-linear differential equation-based stochastic DCM was used to estimate probabilities of these three models. Random effects-based BSM (RFX-BSM) method was used to account for the heterogeneity of model structure across subjects in each group. RFX-BMS provides the expected posterior model probability, which measures how likely a specific model generates the data of randomly selected subjects, and the exceedance model probability, which measures how one model is more likely than any other models, in the group data (Stephan et al., 2010).

### 2.5. Statistical analysis

Schizophrenia and control populations were compared for their age and probabilities of three models using Mann-Whitney U test. Sex was compared between groups using a chi-square test. All data were analyzed using MATLAB (Natick, MA, USA;https://www.mathworks.com). Age and sex covariates were corrected within group and between group comparisons. To account for multiple comparisons, false discovery rate (FDR) correction was applied and a p-value less than 0.05 was considered statistically significant.

## 3. Results

Schizophrenia (n = 67) and healthy control (n = 81) populations were not significantly different in their age (p = 0.99) and sex (p =0.14). Mean age (standard deviation) of schizophrenia patients was 38.46 years (13.96) and that of healthy controls was 37.99 years (11.86). Proportion of female participants was 22.39% (n = 15) in schizophrenia and 25.93% (n = 21) in healthy controls. **Figure 1** shows the results of constrained ICA for DMN, SN, and CEN components in the healthy control group.**Figure 2** displays the results of constrained ICA for DMN, SN, and CEN components in the schizophrenia group.

**Figure 1:**
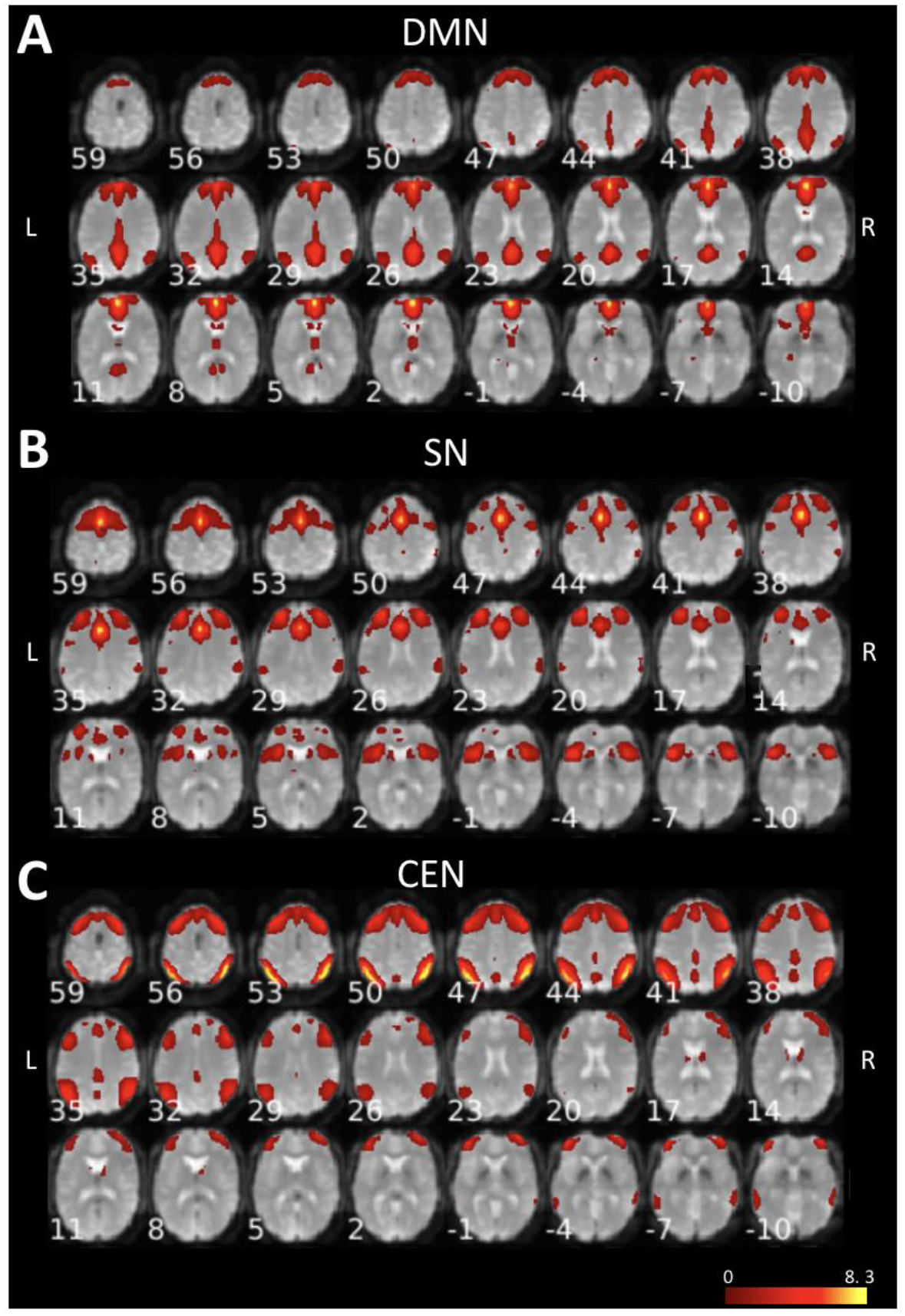
T-value maps of **A**) DMN, **B**) SN, and **C**) CEN from constrained ICA overlaid on mean BOLD images in healthy control (CN) group. Axial view of the brain slices with corresponding z-coordinates are shown (L: Left; R: Right).

**Figure 2:**
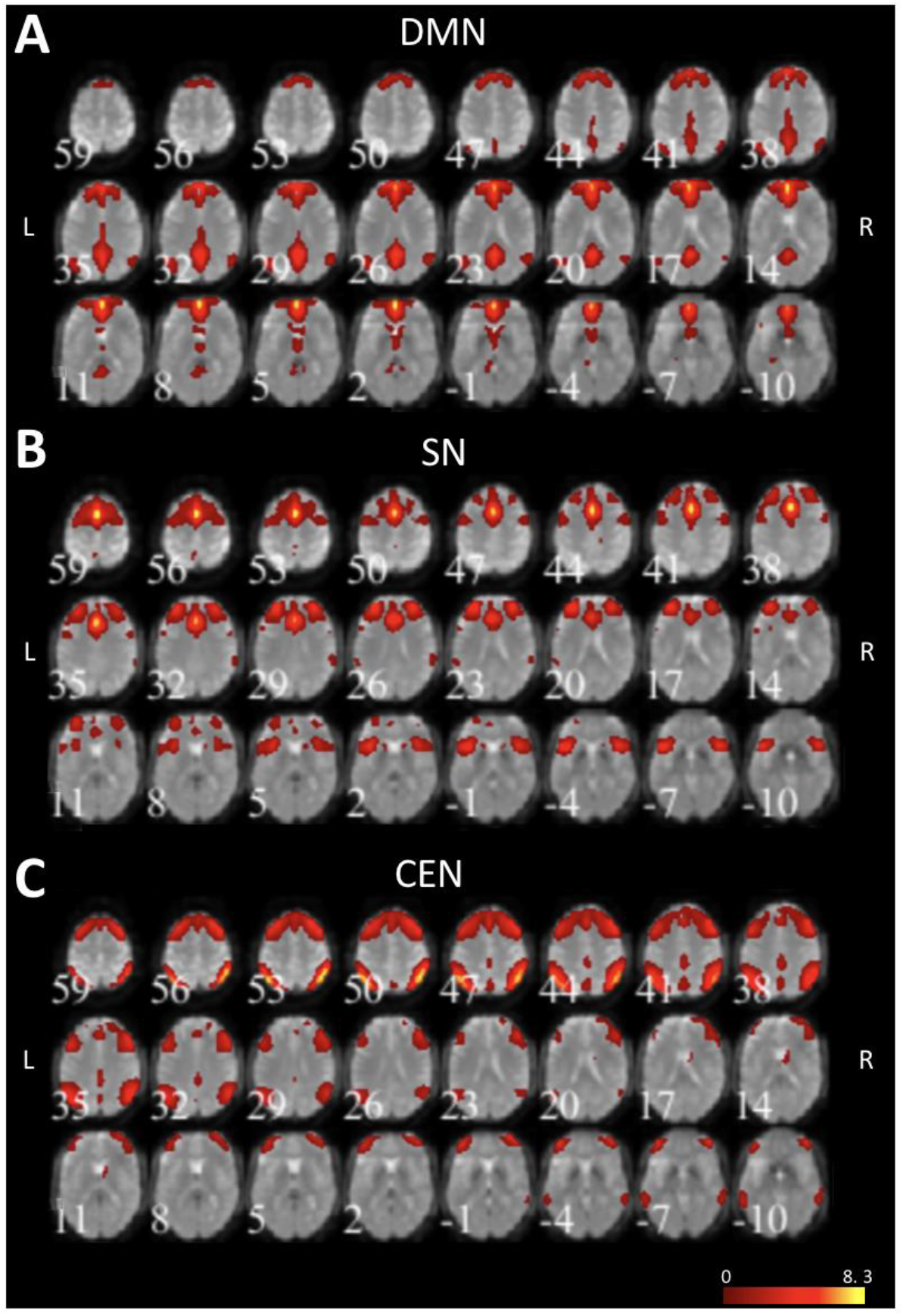
T-value maps of **A**) DMN, **B**) SN, and **C**) CEN from constrained ICA overlaid on mean BOLD images in schizophrenia (SZ) group. Axial view of the brain slices with corresponding z-coordinates are shown (L: Left; R: Right).

The model architectures among DMN, SN, and CEN are illustrated in **Figure 3**. In model 1 (Figure 3A), DMN modulates the reciprocal connections between SN and CEN. In model 2 (Figure 3B), SN modulates the reciprocal connections between DMN and CEN suggesting the controlling/coordinating role of SN. In model 3 (Figure 3C), CEN modulates the reciprocal connection between DMN and SN. DCM RFX-BMS analyses were performed in schizophrenia and healthy control groups (**Figure 4**). The first column (A, C) shows the expected posterior model probability and exceedance probability in the healthy control group. The second column (B, D) shows the expected posterior probability and exceedance probability in the schizophrenia group. The RFX-BMS analyses revealed that model 2 had a higher probability value than the other two models in healthy controls. In the schizophrenia group, model 2 did not have a higher probability while model 3 had a noticeable probability value.

**Figure 3:**
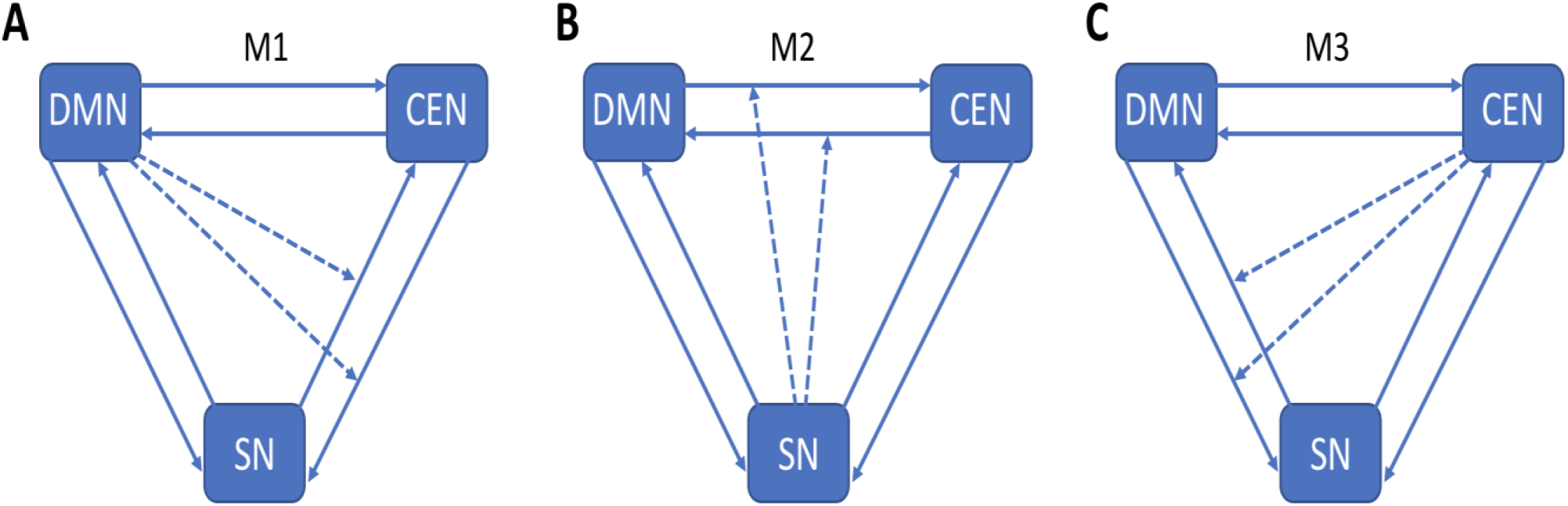
Model architectures: Full intrinsic connections among DMN, SN, and CEN. **A**) Model 1 (M1) specifies nonlinear modulations of DMN on the reciprocal connections between SN and CEN, B) Model 2 (M2) specifies nonlinear modulations of SN on the reciprocal connections between DMN and CEN, and **C**) Model 3 (M3) specifies nonlinear modulations of CEN on the reciprocal connections between DMN and SN.

**Figure 4:**
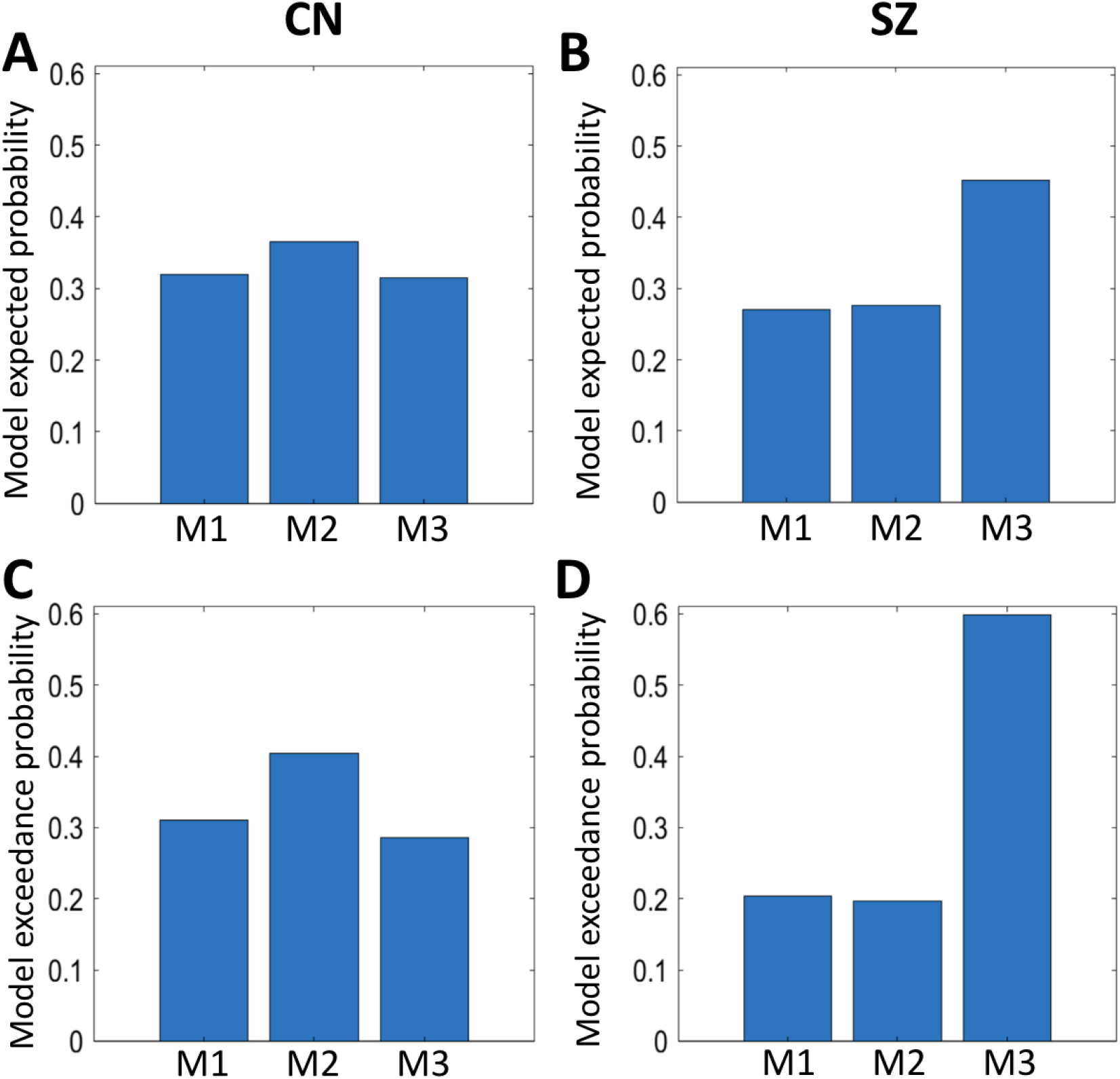
Expected posterior probability and exceedance probability: First column (**A, C**) displays results in healthy controls (CN) indicating that model 2 (M2) has higher probability value compared to other models (M1 and M3). Second column (**B, D**) displays results in schizophrenia (SZ) group suggesting that M2 has lower probability value compared to model 3 (M3).

These three models were further compared within the healthy control and schizophrenia groups (**Figure 5**). In the healthy control group (Figure 5A), we found that model 2 has significantly higher probability compared to model 1 (p < 10^−8^; Mann-Whitney U test) and model 3 (p < 10^−12^) while probabilities of model 1 and model 3 were similar (p = 0.05). In the schizophrenia group, we revealed that model 3 has significantly higher probability compared to model 1 (p < 10^−16^) and model 2 (p < 10^−16^) while probabilities of model 1 and model 2 were not significantly different (p > 0.05). These three models were further compared between the healthy control and schizophrenia groups. This comparison demonstrated significant group differences in model 1 (p < 10^−9^), model 2 (p < 10^−13^), and model 3 (p < 10^−18^) (**Table 1**).

**Figure 5:**
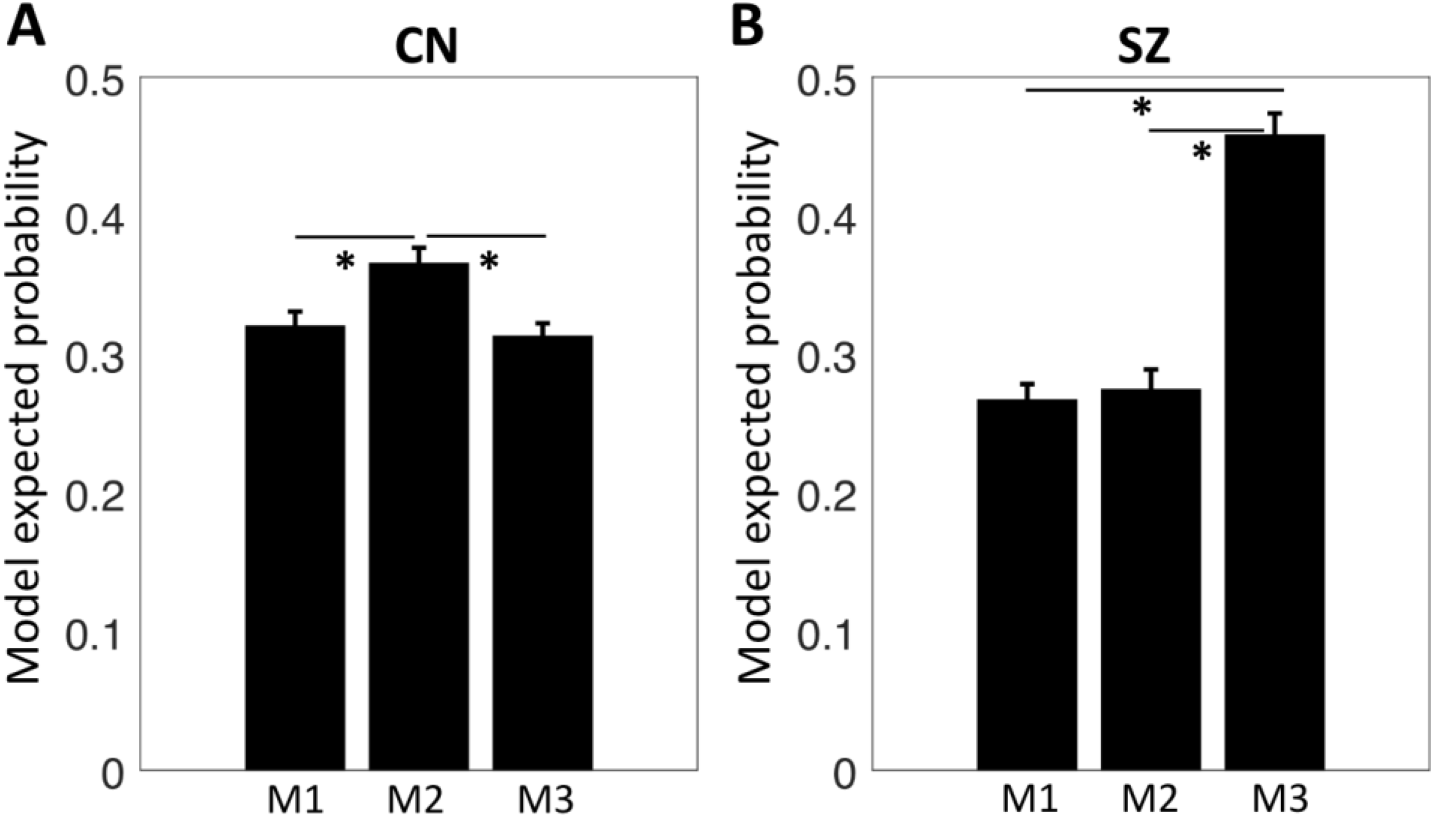
Probability comparison among models within each group: **A**) significantly higher probability value of model 2 (M2) compared to other models within healthy control (CN) group, and **B**) significantly lower probability value of model 2 (M2) compared to model 3 (M3) within schizophrenia (SZ) group. (* indicates FDR-corrected p < 0.05; error bars represent the standard error of mean over participants in each group).

**Table 1:** Model probability value comparison between healthy control (CN) and schizophrenia (SZ) groups. (SD: standard deviation).

| Model | CN<br>Mean (SD) | SZ<br>Mean (SD) | CN vs. SZ<br>P-value |
| --- | --- | --- | --- |
| M1 | 0.32 (0.09) | 0.27 (0.09) | $3.10 \times 10^{-10}$ |
| M2 | 0.37 (0.10) | 0.27 (0.12) | $7.47 \times 10^{-14}$ |
| M3 | 0.31 (0.08) | 0.46 (0.13) | $3.61 \times 10^{-19}$ |

## 4. Discussion

This study tested the hypothesis that SN regulates DMN and CEN in healthy controls and whether such a controlling mechanism is altered in schizophrenia. We showed that SN-based controlling mechanism was maintained in healthy controls while this was disrupted in the schizophrenia group. Adequate interactions between the networks have been argued to be important for effective maintenance of normal cognitive states (Bressler and Menon, 2010; Chand and Dhamala, 2016a; Menon, 2015; Uddin, 2015). Prior investigations have consistently demonstrated that SN drives DMN and CEN during both resting state and tasks in healthy younger and older populations (Chand and Dhamala, 2016a; Chand et al., 2017b; Goulden et al., 2014; Sridharan et al., 2008). Our current results in healthy controls have replicated those reports in existing studies. Our findings further demonstrated that there is disruption of SN-based control mechanism on DMN and CEN in schizophrenia. We tested this controlling mechanism by implementing nonlinear modulations (Chand et al., 2017a, b; Goulden et al., 2014; Li et al., 2011; Stephan et al., 2008) in fully connected intrinsic interactions among DMN, SN, and CEN.

SN-based interactions have been previously described in terms of its distinctive cyto-architecture. SN nodes consist of a special type of neurons named VENs, which are relatively larger in size than other neurons such as pyramidal neurons. These VENs facilitate the dynamic coordinated activity of SN with other networks, including DMN and CEN (Allman et al., 2005; Sridharan et al., 2008). Anterior insula—a key node of SN—connects functionally to CEN (Vincent et al., 2008) and has direct white-matter connections to other areas, including the anterior cingulate cortex (Bonnelle et al., 2012; Jilka et al., 2014), inferior parietal lobe (Uddin et al., 2010), and temporo-parietal junction (Kucyi et al., 2012). Such connections of the anterior insula with prefrontal, precentral operculum, parietal, and temporal cortices are widely replicated in monkeys and humans (Cerliani et al., 2012; Mesulam and Mufson, 1982a, b; Mufson and Mesulam, 1982). Those rich structural and functional connections might support anterior insula to coordinate activity in many cognitive processes, including reorientation of attention (Ullsperger et al., 2010) and switching between cognitive resources (Uddin and Menon, 2009). Another key node of SN—anterior cingulate cortex—is primarily involved in the increased cognitive control and conflict monitoring (Egner, 2009) and it regulates neural activities in association with anterior insula during difficult tasks (Chand and Dhamala, 2016a). Thus, the control mechanism of SN nodes—anterior insula and anterior cingulate cortex—has been argued to be central for active and passive cognitive states in healthy individuals. In schizophrenia, disruption of such coordinated activities of SN might be associated with the gray matter reductions in SN nodes (Gupta et al., 2015) and the impairment of highly sensitive VENs and white-matter fiber tracts (Allman et al., 2010; Allman et al., 2005; Bonnelle et al., 2012; Butti et al., 2013; Sridharan et al., 2008; Watson et al., 2006).

Past studies show that DMN activities alter in schizophrenia (Sendi et al., 2021). Decreased modulations of DMN in schizophrenia compared to healthy controls in our study might indicate such alterations. CEN, especially DLPFC, has rich connections with many brain regions, such as visual, somatosensory, and auditory areas. DLPFC receives neural information from occipital, parietal, and temporal cortices, and it is important in a wide range of cognitive processes, including executive functions (Chand et al., 2016; Miller and Cohen, 2001; Petrides and Pandya, 1999). There are conflicting previous reports on the role of CEN in schizophrenia and other brain diseases, especially whether DLPFC region hypo-activates or hyper-activates remains poorly understood (Chand et al., 2017c; Diener et al., 2012; Lewis, 2012; Potkin et al., 2009). Our results suggest that CEN modulation (model 3) is greater in schizophrenia compared to healthy controls implying hyper-modulations of CEN. Overall, these findings and existing literature taken together indicate that SN-based modulation on DMN and CEN is achieved in healthy individuals, however this controlling mechanism is disrupted in schizophrenia.

Although we leveraged a novel network approach by implementing the nonlinear modulations among DMN, SN, and CEN, and uncovered the disrupted SN-based controlling mechanism in schizophrenia, it is worth noting the following limitations. First, the sample size is not relatively large and replication of these findings in larger samples is recommended. Second, schizophrenia subjects included in this study are chronic patients and future study in non-medicated first-episode schizophrenia will be crucial to understand the early markers of SN-based controlling mechanisms. Third, there is an urgent need of employing the unified network approach to understand other neural networks and future studies should focus in that direction. Lastly, recent findings show that schizophrenia patients exhibit heterogeneity in other neuroimaging modality, especially in neuroanatomical volumetric measurement (Chand et al., 2020). To test whether such heterogeneity exists in the functional connectivity patterns among neural networks, a significantly larger sample size will be required in the future.

In summary, this study evaluated the resting state functional interactions among DMN, SN, and CEN in healthy controls and schizophrenia. SN-based modulation on DMN and CEN was present in healthy individuals, however such SN-based controlling mechanism was disrupted in schizophrenia. These findings shed light on our current understanding of how DMN, SN, and CEN coordinate among each other in schizophrenia.

## Author contributions

G.B.C. conceived the study, analyzed imaging and demographic data, and supervised the study. G.B.C., D.S.T., and B.S. participated in the conceptualization, drafted the manuscript, critically reviewed and approved the final version of manuscript.

## Acknowledgements

Authors acknowledge the publicly available dataset. Data used in preparation of this article were obtained from the SchizConnect database (http://schizconnect.org). As such, the investigators within SchizConnect contributed to the design and implementation of SchizConnect and/or provided data but did not participate in analysis or writing of this report. Data collection and sharing was funded by NIMH cooperative agreement 1U01 MH097435. Data was downloaded from the COllaborative Informatics and Neuroimaging Suite Data Exchange tool (COINS; http://coins.mrn.org/dx) and data collection was performed at the Mind Research Network, and funded by a Center of Biomedical Research Excellence (COBRE) grant 5P20RR021938/P20GM103472 from the NIH (Aine et al., 2017; Cetin et al., 2014).

